# Sex-specific association between urinary kisspeptin and pubertal development

**DOI:** 10.1101/2021.07.12.452029

**Authors:** Rafaella Sales de Freitas, Thiago Fonseca Alves França, Sabine Pompeia

## Abstract

Kisspeptins are critical neuropeptides for puberty onset and progression, playing a pivotal role in the reactivation of the hypothalamus-pituitary-gonadal axis in late childhood. Despite their importance, little is known as to how kisspeptin peripheral concentrations are related to sexual maturation in humans, specially using non-invasive measures that allow more widespread testing. Using a cross-sectional design, we investigated whether peripheral kisspeptin concentrations, measured by enzyme-linked immunosorbent assays (ELISA) in two-hour retention midstream urine, are associated with developmental markers in 209 (120 girls) typically developing, 9 to 15-year-olds. Developmental variables were age, self-reported pubertal status using the Pubertal Development Scale, and saliva concentrations of hormones that indicate gonadal (testosterone) and adrenal (dehydroepiandrosterone sulfate) functioning. We found marked sexual differences in urine kisspeptins (controlled for body mass index and socioeconomic status). While concentrations were similar in both sexes until the age of around 12 years, in males there was a positive linear correlation with all developmental measures thereafter, while in girls, kisspeptin concentrations did not change. Of note, our results are in line with those of previous studies using more invasive methods (e.g. blood samples), which indicates that kisspeptin concentrations from two-hour retention midstream urine have potential in exploring sex-specific peripheral action of these peptides.

---

Adolescence is a period of life that involves social, cognitive, psycho-emotional and physical/pubertal changes (Worthman et al., 2019). The physiological aspects of puberty revolve around a set of marked endocrine changes that reflect maturation of the gonads, acceleration of height growth and development of secondary sexual characteristics. The kisspeptin peptides, produced by neurons in two hypothalamic nuclei (the arcuate nucleus and the anteroventral periventricular nucleus), also play a key role in these changes. Kisspeptins act on their receptors (KISS1R, also known as G coupled protein receptor 54 [GPR54]) located on gonadotropin releasing hormone (GnRH)-neurons in the hypothalamus. This increases GnRH-neurons’ pulsatile secretion of GnRH and is essential for the onset of puberty and regulation of fertility (Clarke et al., 2015; Smith & Clarke, 2007; Oakley et al., 2009). Increases in GnRH stimulate the pituitary glands to release gonadotropins (luteinizing hormone [LH] and follicle-stimulating hormone [FSH]), also in a pulsatile manner, leading to the production of sex steroids by the gonads (e.g. testosterone by the testis; progesterone and estrogens by the ovaries) (Clarke et al., 2015; Smith & Clarke, 2007; Oakley et al., 2009). The elements involved in the control of the hypothalamic-pituitary-gonadal (HPG) axis are highly conserved across vertebrates, including the kisspeptin gene (KISS1; Tsutsui et al., 2010), which in humans encodes a precursor protein that can be cleaved into three shorter peptides (kisspeptins 13, 14 and 54), all of which can activate the KISS1R receptors (Kotani et al., 2001).

In addition to the compelling evidence regarding the direct action of kisspeptins on the hypothalamus, there is also growing evidence that kisspeptins activate non-GnRH cells, both in the brain and elsewhere in the body, participating in the regulation of both reproductive and metabolic systems (for recent reviews, see Cao et al., 2019; Dudek et al., 2018; Uenoyama et al., 2016). Indeed, circulating concentrations of kisspeptins are known to change under other conditions related to reproductive functioning beyond puberty, such as during pregnancy (Jayasena et al., 2014) and different phases of the menstrual cycle (Zhai et al., 2017). Furthermore, kisspeptins and their receptors are implicated in socioemotional behavior, modulating anxiety, mood, and fear, which may affect reproductive behavior (Mills et al., 2018). Transcription of genes encoding kisspeptins and their receptor are also found in many human tissues, such as the testis and ovaries, placenta, heart, liver and skeletal muscles (reviewed in Clarke et al., 2015; Wolfe and Hussain, 2018), suggesting widespread action of kisspeptins throughout the body. Importantly, peripheral administration of kisspeptins increases circulating gonadotrophins in humans (Thomson et al., 2004; Clarke et al., 2015), suggesting that peripheral concentrations of these peptides can affect the brain. Therefore, quantifying peripheral, circulating kisspeptin concentrations may be help elucidate their role in human development during adolescence.

Research on kisspeptins may well improve the characterization of pubertal trajectory because there are as yet still no objective measures that represent all stages of puberty, resulting in limitations in our understanding of the physiology of sexual development. Puberty is a complex, multisystemic process that effects health and well-being and is influenced not only by hormonal changes, but also by genetic, metabolic/nutritional, socioemotional and socioeconomic factors (Worthman et al., 2019). However, despite the growing interest in kisspeptins and the flood of new research on these peptides, their effects on pubertal onset and progression in humans are still little understood, with available evidence coming mainly from the study of circulating kisspeptins on clinical populations and in non-human animals (Worthman et al., 2019).

An important issue that must be considered when exploring kisspeptin effects is the sexual dimorphism in the distribution of its receptors, its expression, and in serum kisspeptin concentrations, which may explain developmental differences between males and females (see Bianco, 2012; Kaufman, 2010). Additionally, activation of the HPG axis, which leads to gonadal-dependent development of secondary sexual characteristics (such as breast and genital growth), is not the only change that occurs during puberty (Dorn et al., 2006). Puberty also involves alterations in the adrenal glands, independently of gonadal axis alterations (Dorn et al., 2006; Havelock et al., 2004). These adrenal changes contribute to the development of other secondary sexual characteristics, such as growth of body and pubic hair and skin changes that result mainly from increases in adrenal androgen concentrations, especially dehydroepiandrosterone (DHEA) and its sulphate (DHEA-S) (Dorn et al., 2006; Havelock et al., 2004). There is no evidence that these adrenal changes are related to kisspeptins, but this issue has not been explored in the literature.

Regarding changes in serum/plasma kisspeptins during puberty, there is also a considerable amount of controversy. Some studies have shown that, in girls, kisspeptin concentrations measured in blood increase from pre-puberty to the initial stages of puberty, but not from then on (Jayasena et al., 2013; Xiaoyu et al., 2011; Zhu et al., 2016). Girls with idiopathic central precocious puberty also present higher serum concentrations of kisspeptins than normally developing controls (Yang et al., 2015), further suggesting a role of peripheral kisspeptins in initiating puberty in this sex. In contrast, Zhu et al. (2016) found a marked increase in serum concentrations from the first stages of puberty in boys, but not in girls, in whom concentrations were much lower and more stable after pubertal onset. Conversely, Pita et al. (2011) found no changes in plasma kisspeptins from prepubertal to pubertal youngsters, either boys or girls. There also seems to be a positive linear association of peripheral kisspeptins and androgen concentrations that stimulate the development of secondary sexual characteristics in both sexes, such as testosterone and DHEA-S (Yilmaz et al., 2014; Zhu et al., 2016), but these studies did not compare these correlations between males and females, so it is unresolved whether there are sex differences in these associations. Therefore, the association of peripheral kisspeptin concentrations with pubertal development in each sex clearly needs clarification.

In humans, kisspeptin concentrations can be quantified not only in the blood (plasma/ serum) (Kataguiri et al., 2007), but also in urine and saliva, although saliva samples are much less able to detect kisspeptine changes (Jayasena et al. 2014). Urine kisspeptin detection is possible because these peptides consist of small molecules that pass through the glomerular filtration membrane in the kidneys (Zhai et al. 2017). However, while serum and urine measures of kisspeptin present the same pattern of variation, measurements made in urine samples are less sensitive to variations in peripheral concentrations of kisspeptins (Jayasena et al. 2014). Still, urine collection is a non-invasive method, which is a considerable advantage compared to obtaining blood samples, especially when working with vulnerable populations, such as children and adolescents, as well as pregnant women. Moreover, urine samples are easier to collect than blood samples, the later requiring trained staff to draw blood.

Despite demonstrating that urine could be a good, non-invasive method for measuring kisspeptins, Jayasena et al. (2014) and Zhai et al. (2017) did not provide detailed information about sample collections and analyses. For example, what was the urine retention time? Was midstream urine used? Also, urine essays used in these studies were carried out in pregnant women (Jayasena et al., 2014) and women of reproductive age over the menstrual cycle (Zhai et al., 2017), so it is unknown if this would be a sensitive method to detect changes in peripheral kisspeptin concentrations during pubertal development.

Our objective in this study was to test whether urine detection of kisspeptins in typically developing early male and female adolescents relates to classic developmental markers such as chronological age and pubertal status [as assessed by self-reported pubertal stage using the Pubertal Developmental Scale (PDS), and salivary concentrations of hormones related to gonadal [testosterone] and adrenal (DHEA and DHEA-S) development], which change during the pubertal trajectory in both sexes (e.g. Shirtcliff et al., 2009). We controlled our analyses for body mass index (BMI) because kisspeptins also regulate glucose homeostasis, feeding behavior and body composition, variables that are related to pubertal development (see Bianco, 2012; Pita et al., 2011; Zhu et al., 2016). Socioeconomic status (SES) is also a factor that alters pubertal timing (see Worthman et al., 2019), so we controlled for this as well. Due to lack of information on urine samples used in prior studies that assessed urine kisspeptins, we collected midstream urine with a minimum of two-hour retention in order to establish if this is an adequate method to measure changes in kisspeptin concentrations related to pubertal development.

## Methods

### Participants

Our convenience sample was composed of 209 (120 females) healthy, native Portuguese speaking, 9- to 15-year-old youngsters, – an age interval spanning the full range of pubertal development stages in most adolescents (Dorn et al., 2006). Exclusion criteria, assessed via reports given by guardians, were: (1) use of chronic medication (chosen as a criterion to rule out the presence of clinical disorders), (2) use of drugs that could impact pubertal development and metabolism, and (3) having been held back in school for a year or more and/or being a student with special needs (both of which could indicate the presence of clinical and/or cognitive issues).

### Procedure

The study protocol was approved by the Ethics Committee of the University where it took place (# 56284216.7.0000.5505). All experimenters were trained before data collection. Participants were only included after obtaining signed informed consent from their guardians and their own assent, as per local ethical guidelines. Participants were recruited from public and private schools in the city of São Paulo, in the Southeast region of Brazil and tested at their own school. Their guardians provided demographic and health information. Each participant was invited to participate in two sessions. The first was an individual session in which participants answered questionnaires (including the PDS) and performed a series of cognitive tasks which will be discussed elsewhere. The second session happened in groups, and its purpose was to obtain saliva samples (used for measuring testosterone and DHEA-S) and others biomarkers that are not discussed here. The saliva samples were free of visual blood traces, and were collected by passive drool 10 min after a light mouthwash with water. Anthropometric measurements and urine samples were also obtained during the second session.

Brazilian students study either in the morning, beginning at around 07:00h, or in the afternoon, starting around 12:30h, so it was not possible to obtain all samples at the same time of day. In order to control for circadian variations, we established that samples would be collected either between 10:00-12:00h or between 13:00-16:00h to coincide with each participant’ school shift. A “science partner” certificate was awarded to all participants for taking part in the study and their travel expenses were reimbursed.

### Measures

#### Urinary kisspeptin concentrations

Midstream urine was obtained after at least two-hour retention periods. They were collected as follows: participants were instructed to go to the toilet, dispense a little urine (first pass) to avoid bacterial contamination, and then fill the collector with about 10 ml of urine. Collected biological samples were transported in ice-cooled Styrofoam boxes and aliquots were stored in a freezer at −80 °C until analyzed.. Concentrations of kisspeptins were determined by enzyme-linked immunosorbent essays (ELISA) for human samples following the supplier’s guidelines, in uniplicate (KIT Cloud-Clone Corp. [CCC, Wuhan]). The kit used antibodies produced against the full human KISS1 protein, minus the signal peptide. In other words, it used the sequence from Glu20 to Gly138. This sequence encompasses kisspetins 54 and its shorter forms.

Urine samples of approximately 10 ml were obtained and transported in ice-cooled Styrofoam boxes until they were centrifuged at 1,000 x g for 20 min, after which the supernatant was aliquoted at 1200 μl. Aliquots were stored in a freezer at −80°C until thel. Aliquots were stored in a freezer at −80°C until the analyses. Detection range is 31.2-2,000 pg/mL, with minimum detectable dose of 13.1 pg/mL. Intra- and inter-assay coefficients of variance are < 10% and < 12%, respectively.

#### Other developmental markers

Participants’ age was considered in number of months.

Pubertal status was self-assessed using the PDS, adapted from Carskadon and Acebo (1993) for local use by Pompeia et al. (2019). The scale contains a series of questions regarding the development of secondary sexual characteristics, assessing characteristics related to both gonadal and adrenal factors: body hair development, growth spurt and skin changes for participants of both sexes, voice changes and facial hair growth for boys, and breast growth in girls. All these were measured using a 4-point Likert scale (ranging from 1=“not yet started” to 4=“seems complete”; with 0=“I don’t know”, coded as missing data). Girls were also asked to indicate whether they already had their first menstruation (menarche, score as 1 = no, or 4 = yes). The final scores were the mean ratings in all five sex-specific questions (Carskadon and Acebo, 1993).

Hormonal measures included free salivary testosterone and DHEA-S concentrations. For these measures, the participants’ passive saliva was collected 10 min after a light mouthwash as described by the essay kit (Salimetrics, State College, PA). Specific instructions required participants to think of their favorite foods, expect saliva to accumulate in their mouth and then to tilt their heads down and drop the saliva in the collector without forcing or producing sputum. The collectors were then inspected and samples contaminated with blood were discarded and other samples were obtained. Of note, participants were instructed a day before the second session to avoid eating, brushing their teeth and chewing gum one hour before sample collections, because this could interfere with saliva production and contaminate them with blood.

Hormone concentrations were determined from saliva aliquots (700 μl) using ELISA, in uniplicate, following the supplier’s guidelines. Absorbance was measured using a spectrophotometer reader Stat Fax model 2100 from Awareness Technology. The concentrations were calculated in relation to standard curves and expressed in pg/ml for testosterone and ng/ml for DHEA-S. The test sensitivities were 0.94 pg/mL and 0.05 ng/mL, respectively. Some random samples were tested in duplicates. In these, the mean coefficients of variation were 4.4% for testosterone samples and 4.1% for DHEA-S.

#### Other control measures

Participants’ SES was assessed using a measure that is adequate considering local/cultural specificities, for example, rampant unemployment and informal occupations (see Colom and Flores-Mendoza, 2007): family purchasing power in accordance with the guidelines of the Brazilian Association of Market Research (see http://www.abep.org.br; for a version in English, see http://www.abep.org/Servicos/Download.aspx?id=11). This questionnaire, answered by one of the guardians, attributes points based on the number of items present at the respondents’ homes (e.g., number of cars, motorcycles, bathrooms, refrigerators, freezers, computers, DVDs, washing and drying machines, dishwasher, microwave, presence of a housemaid who works on all days in a week), whether the street where they live is paved or not, if the house has or does not have piped water, and the mean educational attainment of the parents/guardians (here we deviated from the scale, which considered the education of the breadwinner of the household, because many families expressed difficulty determining who was the breadwinner). We used the points obtained from this scale, divided by the number of residents in the home (*per capita* SES), as a continuous variable in the statistical analyses.

Body Mass Index (BMI: kg/m^2^) was calculated using weight, determined after removal of coats and shoes with an OMRON HBF-514C Body Control scale (to the nearest 0.1 kg), and standing height (to the nearest cm), recorded in bare feet using a stadiometer. Both weight and height were measured twice and the average values were used.

#### Statistical analyses

The descriptive and inferential analyses were performed with the SPSS, v. 21. Pearson correlation indices were used to explore the relations between kisspeptin concentrations and markers of development (age, PDS, testosterone and DHEA-S). Using univariate General Linear Models (GLM), we investigated how the urinary concentrations of kisspeptins (treated as a dependent variable) related with developmental indicators (each in a separate model), controlling for sex (modeled to interact with the developmental markers), SES and BMI, which are known to influence pubertal trajectories (Bianco, 2012; Pita et al., 2011; Zhu et al., 2016). These models generate parameters common to ANOVAs (*df*, *F* and *p* values) and regressions (unstandardized *B* regression coefficients; adjusted *R*^*2*^ to allow comparisons of models; and multiple *R*^*2*^, which indicates the proportion of variance explained by the models). Multiple *R*^*2*^ values from .13 to .25 were considered medium effect sizes, and large effect sizes when above .26 (Ellis, 2010; Kotrlik et al., 2011). The level of significance adopted for all the inferential analyses was *p* ≤ .05. There was no imputation of missing data, nor removal of outliers.

## Results

The sample represented participants in all different stages of sexual maturation, varying ages, SES and BMI (Table 1). These variables were all regressed on the self-assessed PDS scores, and the resultant intercepts and unstandardized regression coefficients (*B*, with their respective 95% Confidence Interval [CI]) are also reported in Table 1 so that the reader can gauge the values per sexual maturation status (PDS scores). There were some missing data [4 (1 for girls and 3 for boys) for DHEA-S, 3 (1 for girls and 2 for boys) for testosterone and 4 (2 for girls and 2 for boys) for BMI]. The databank can be found at https://osf.io/dqesj/.

**Table 1.** Descriptive data [mean, standard deviation (*SD*), Minimum (*Min*), Maximum (*Max*)] of all analyzed variables included in this study, by sex, and Intercept (*Interc.*) and unstandardized regression coefficient (*B*) [with respective 95% Confidence Interval (*CI*)] for the regression of each variable on scores in the self-assessed Pubertal Development Scale (PDS).

| Variable | Girls (N=120) |  |  |  |  |  |  |  |  |  | Boys (N=89) |  |  |  |  |  |  |  |  |  |
| --- | --- | --- | --- | --- | --- | --- | --- | --- | --- | --- | --- | --- | --- | --- | --- | --- | --- | --- | --- | --- |
|  | Mean | SD | Min | Max | Interc. | ±95% CI |  | B | 95% CI |  | Mean | SD | Min | Max | Interc. | ±95% CI |  | B | B ±95% CI |  |
| **PDS (escore)** | 2.7 | 0.7 | 1.2 | 3.8 |  |  |  |  |  |  | 2.1 | 0.6 | 1.0 | 3.6 |  |  |  |  |  |  |
| **Age (years)** | 12.6 | 1.7 | 9.0 | 16.0 | 8.03 | 7.20 | 8.86 | 1.69 | 1.41 | 1.98 | 12.2 | 1.7 | 9.0 | 15.0 | 7.99 | 7.11 | 8.87 | 2.00 | 1.62 | 2.39 |
| **Kisspeptin (pg/ml)** | 47.29 | 7.72 | 32.93 | 72.03 | 48.131 | 42.357 | 53.905 | -0.311 | -2.381 | 1.758 | 78.69 | 34.77 | 32.79 | 211.08 | 18.733 | -3.130 | 40.596 | 28.401 | 18.482 | 38.320 |
| **Testosterone (pg/ml)** | 2.66 | 1.85 | 0.36 | 11.58 | 11.126 | -2.256 | 24.509 | 6.497 | 1.779 | 11.214 | 2.89 | 1.96 | 0.12 | 11.28 | -48.494 | -77.592 | -19.396 | 48.289 | 35.041 | 61.536 |
| **DHEA-S (ng/ml)** | 29.77 | 20.17 | 4.18 | 112.83 | 0.709 | -0.509 | 1.928 | 0.694 | 0.265 | 1.124 | 50.93 | 46.87 | 3.13 | 254.12 | 0.905 | -0.461 | 2.271 | 0.965 | 0.339 | 1.591 |
| **BMI (kg/m<sup>2</sup>)** | 20.9 | 3.8 | 14.5 | 31.2 | 16.34 | 13.57 | 19.10 | 1.78 | 0.80 | 2.75 | 21.1 | 4.2 | 13.7 | 32.8 | 17.86 | 14.93 | 20.79 | 1.48 | 0.16 | 2.80 |

Note: DHEA-S= dehydroepiandrosterone sulfate; BDI=body mass index.

Pearson correlations of kisspeptin concentrations with the developmental markers were all positive and statistically significant (*p* < .01) for boys: for age, *r* = .624; for PDS, *r* = .595; for DHEA-S, *r* = .277; for testosterone, *r* = .350. None of the correlations were significant for girls (coefficient values ranged from *r* = −.027 to .132).

Table 2 shows the results of the GLMs used to investigate how kisspeptin concentrations (dependent variable) varied according to developmental variables in separate models (age in months, PDS scores, DHEA-S and testosterone).

**Table 2.** Results of the General linear models (GLM) used to determine urine kisspeptin concentrations across development (chronological age and pubertal indicators), per sex, controlled for body mass index and socioeconomic status.

| **Models (df)** | **Predictors** | **F** | **p** | **$\eta^2$** | **B** | **Multiple R<sup>2</sup> (adjusted R<sup>2</sup>)** |
| --- | --- | --- | --- | --- | --- | --- |
| 1 (1,204) | **Age** (months) | 64.61 | <0.001 | 0.245 | 1.126 | 0.56 (0.55) |
|  | Sex (males as a reference) | 44.32 | <0.001 | 0.182 | 140.437 |  |
|  | Interaction Sex vs. Age | 68.68 | <0.001 | 0.257 | -1.125 |  |
|  | Body Mass Index | 0.83 | 0.36 | 0.004 | -0.319 |  |
|  | Socioeconomic Status | 0.21 | 0.65 | 0.001 | 0.214 |  |
| 2 (1,204) | **PDS** (Score) | 38.945 | 0.001 | 0.164 | 28.956 | 0.49 (0.48) |
|  | Sex (males as a reference) | 7.71 | 0.006 | 0.037 | 30.329 |  |
|  | Interaction Sex vs. PDS | 42.70 | <0.000 | 0.177 | -29.200 |  |
|  | Body Mass Index | 0.29 | 0.59 | 0.001 | -0.203 |  |
|  | Socioeconomic Status | 0.04 | 0.84 | <0.001 | 0.104 |  |
| 3 (1,200) | **DHEA-S** (ng/ml) | 10.09 | 0.002 | 0.049 | 4.782 | 0.36 (0.34) |
|  | Sex (males as a reference) | 10.50 | 0.001 | 0.051 | -18.391 |  |
|  | Interaction Sex vs. DHEA-S | 5.86 | 0.02 | 0.029 | -4.096 |  |
|  | Body Mass Index | 0.28 | 0.59 | 0.001 | -0.220 |  |
|  | Socioeconomic Status | 1.09 | 0.30 | 0.006 | 0.583 |  |
| 4 (1,201) | **Testosterone** (pg/ml) | 3.84 | 0.05 | 0.019 | 0.252 | 0.39 (0.38) |
|  | Sex (males as a reference) | 12.55 | 0.004 | 0.600 | -18.292 |  |
|  | Interaction Sex vs. Testosterone | 5.54 | 0.02 | 0.029 | -0.274 |  |
|  | Body Mass Index | 0.05 | 0.83 | <0.001 | 0.089 |  |
|  | Socioeconomic Status | 0.36 | 0.55 | 0.002 | 0.337 |  |

Note: PDS=scores in the Pubertal Development Scale; DHEA-S =dehydroepiandrosterone sulphate.

[Tables 1 and 2 near here]

All models were able to explain a significant proportion (> 34%) of the variance in urine kisspeptine concentrations. All developmental markers had significant effects (*p* values <.002), except for testosterone, which fell short of significance (*p* = .051). Kisspeptins increased with age (*B* = 1.13 pg/ml per month), PDS scores (*B* = 28.96 pg/ml per point), DHEA-S (*B* = 4.78 pg/ml per ng/ml) and testosterone (*B* = .25 pg/ml per pg/ml), but effect sizes in terms of partial eta squared values only reached medium values (*p*η^2^ > .13) in the first two cases. Sex was a significant predictor in all models (*p* values < .006), with boys having consistently higher kisspeptin concentrations than girls. The interaction of sex with each developmental marker was also significant (*p* values < 0.02), indicating that kisspeptin concentrations were not similar among boys and girls at any ages. However, this interaction reached medium effect sizes only for the models with age and PDS scores. BMI and SES were not significant factors in any of the models.

To facilitate the interpretation of these findings, Figure 1 shows kisspeptin concentrations according to each developmental marker, separately by sex. The top-left plot in Figure 1 shows the relationship between kisspeptin concentrations and age in boys and girls. We can see that there is no longer an overlap of the 95% confidence intervals between kisspeptin levels of girls and boys from just before the age of 12 years (144 months) onward, with boys showing higher concentrations thenceforth, while girls show no perceptible variation in kisspeptin concentrations at any age. Regarding the PDS (top-right plot), boys had higher kisspeptin concentrations than girls from PDS score 2 onward. For DHEA-S and testosterone (bottom-left and -right plots, respectively), sex differences are observed in the full range of results, again with boys presenting higher kisspeptin concentrations. In all cases, kisspeptin levels in girls were unchanged with regards to developmental measures.

[Figure 1 near here]

**Figure 1.**
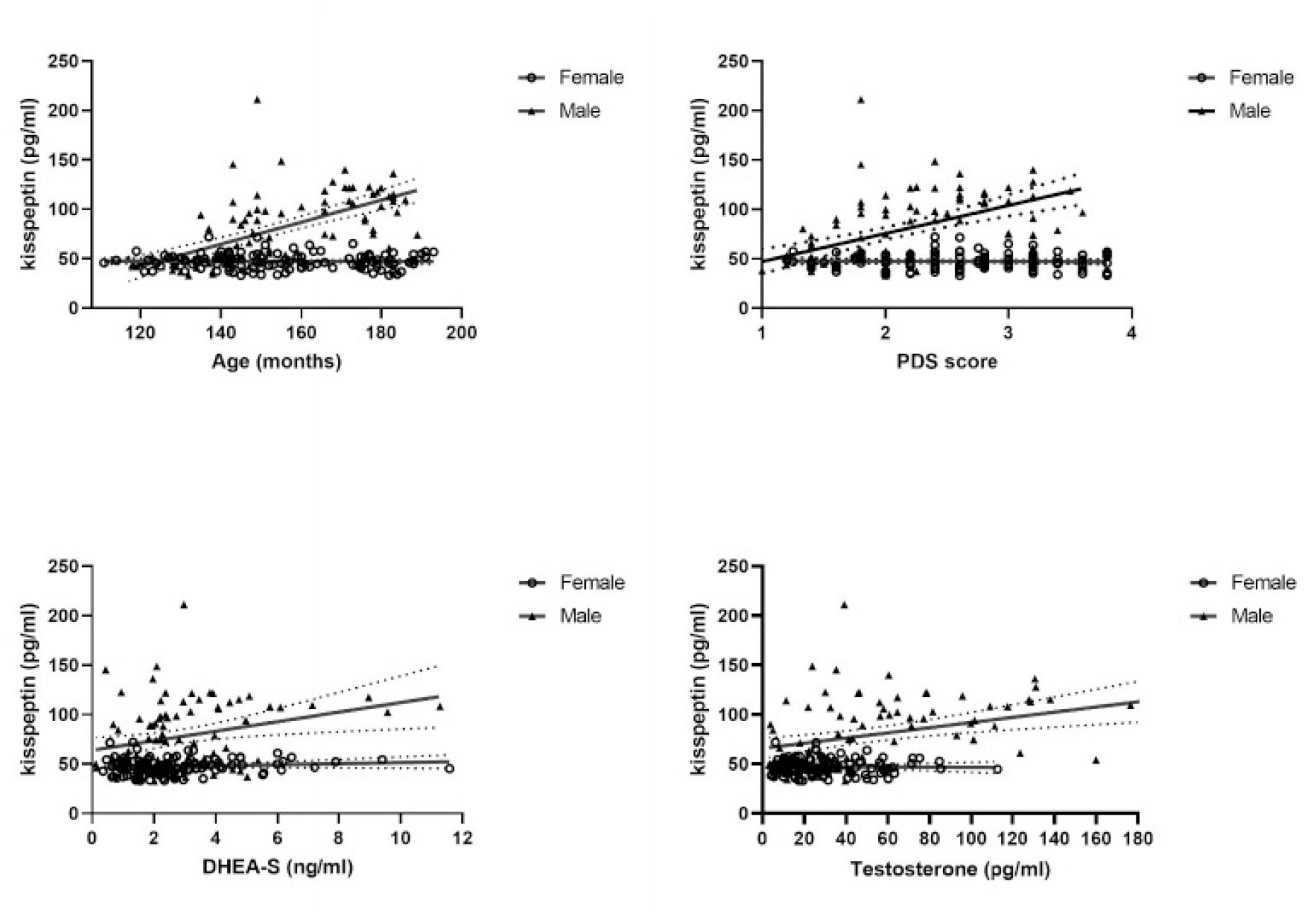
Scatterplots representing individual data of urine kisspeptin concentrations according to developmental markers, per sex. Continuous lines represent simple regressions, and doted lines represent their 95% confidence intervals. This representation is not corrected for socioeconomic status and body mass index, which were non-significant predictors (see main text). PDS=scores in the Pubertal Development Scale; DHEA-S =dehydroepiandrosterone sulphate.

## Discussion

In this exploratory study, we aimed to show whether measuring kisspeptin concentrations from midstream, two-hour urine retention samples was an adequate method to quantify this peptide according to developmental markers (chronological age and pubertal markers) in early adolescents. The results showed that kisspeptin concentrations were very similar in boys and girls in early puberty (PDS scores < 2), at around the ages of 9 to 11 years, but that, thenceforth, these concentrations markedly increased in boys but did not change in girls. These sex differences corroborate the data of Zhu et al. (2016), who measured kisspeptins in the serum of typically developing youngsters. Furthermore, the fact that the study by Zhu et al. (2016) had, like ours, a larger sample size compared to other published studies that measured kisspeptin in blood samples, and used the same commercial kit employed in our study, suggests that urine measurement is a feasible, alternative non-invasive method to quantify peripheral kisspeptins. However, further studies are required to confirm the kits reliability, as discussed below.

The only difference between Zhu et al.’s (2016) and our study was that they found that serum kisspeptin concentrations in girls were higher at the beginning of puberty compared with pre-puberty, confirming the findings of Xiaoyu et al. (2011). This change in kisspeptin levels at the onset of puberty is consistent with the findings of Yang et al. (2015), who showed that girls with idiopathic central precocious puberty had higher serum levels of kisspeptins than normally developing controls, and that, after treatment, these concentrations fell below pre-treatment levels, suggesting a role for peripheral kisspeptin concentrations on pubertal onset in this sex. We did not detect this specific increase in the present study, probably due to the small number of prepubertal girls in our sample – only 26 girls scored less than 2 in the PDS. Furthermore, Zhu et al. (2016) used a discrete marker of pubertal status (Tanner stages), while we used a continuous PDS score that makes this putative effect difficult to show.

Of note, the data from Zhu et al. (2016), Xiaouy et al. (2011), Yong et al. (2015) and ours are not in line with the results of Pita et al. (2011) and Jayasena et al. (2013). Pita et al. (2011) found stable serum kisspeptin concentrations throughout puberty and adulthood, while Jayasena et al. (2013) claimed that plasma concentrations of kisspeptin peaked between the ages of 9 and 12 years in both sexes, after which they decreased until adulthood.

It is difficult to pin down the roots of the disagreement between published studies. One the one hand, there are differences between studies in the methods used to assess kesspeptins. Pita et al. (2011) and Jayasena et al. (2013) quantified kisspeptins using radioimmunoessays with antibodies produced against kisspeptin 54 that where capable of detecting kisspetins 14 and 13/10 and had no cross-reactivity with some known kisspeptin analogues. Our study and the ones from Zhu et al. (2016), Xiaoyu et al. (2011) and Yong et al. (2015) all used ELISA kits. The kit used by Yang et al (2015) employed antibodies produced against human kisspeptin 54, but the manufacturer states they cross-reacted with some of kisspeptins analogues. There is no information about the kit used by Xiaoyu et al. (2011), while the kit employed by Zhu et al. (2016) and by the present study used antibodies produced against the product of the KISS1 gene, which includes the kisspetin 54 sequence, but also several additional aminoacids. This manufacturer (KIT Cloud-Clone Corp. [CCC, Wuhan]) states the kit has high sensitivity and specificity, but there is limited information about cross-reactivity with kisspeptin analogues and about specificity for particular kisspeptins. How the differences in antibodies and detection methods used in the different studies affect the results is unknown and requires further investigation, especially because this variability in kisspeptin measurement methods is ubiquitous in the literature.

On the other hand, while it is tempting to give more weight to the results of Pita et al. (2011) and Jayasena et al. (2013) on the grounds of the apparently lower risk of bias in their kisspeptin measurement methods, it is important to note that this is not the only difference between the studies. For example, in contrast to the relatively large sample size used in the present study and in Zhu et al. (2016), both Pita et al. (2011) and Jayasena et al. (2013) used very small samples – especially considering the analyses by sex and age during puberty. This, and the fact that those studies apparently did not control for multiple comparisons, increases the risk of unreliable results.

As shown here, peripheral kisspeptin concentrations have been found to be positively associated with testosterone and DHEA-S in pubertal boys (see Zhu et al., 2016). However, we failed to find reports of these correlations in pubertal girls. Importantly, the fact that DHEA-S is related to kisspeptins does not necessarily mean that these peptides act on adrenal pubertal systems. Such an association would be expected due to the relative synchronicity between gonadal and adrenal maturation, despite the independent nature of these processes (see Dorn et al., 2006). Although higher concentration of testosterone and DHEA are found in more advanced pubertal stages in both sexes (Shirtcliff et al., 2009), Van Hulle et al. (2015) claim that these hormones do not vary in parallel with changes in secondary sexual characteristics measured by subjective scales, nor seem to share the same mechanisms in males and females (Van Hulle et al., 2015). This can help explain why these hormones were only related to kisspeptins in boys, although they only reached small effect sizes.

Overall, our results, together with those of studies in other pubertal samples, show that increases in peripheral kisspeptins are not only associated with the initiation of puberty, but are also related to sexual maturation trajectory in boys, while girls present much lower concentrations of peripheral kisspeptins and show little developmental variation except from the passage from pre-puberty to early puberty (Xiaoyu et al., 2011; Zhu et al., 2016). These sex differences may seem puzzling at first, but some additional lines of evidence regarding the roles of kisspeptin in development and beyond can shed some light on the meaning of these results. First, there are sex differences in the sensitivity and/or number of central KISS1 receptors (see Bianco, 2012; Kaufman, 2010 Nabi et al., 2018), that can also vary across puberty, at least in boys (Nabi et al., 2018). Second, kisspeptins are also produced by several peripheral tissues that are sex specific, including the testis and ovaries, and may have sex-specific roles in the reproductive systems of males and females that go beyond those related to puberty itself (as reviewed in Cao et al., 2019).

Third, as due to the widespread expression of kisspeptin in the periphery beyond the gonads, including in the adrenals glands, liver, pancreas, adipose tissue and blood vessels (reviewed in Wolfe and Hussain, 2018), at least part of the sex differences in peripheral kisspeptin concentrations may well be explained by other physiological roles of kisspeptins. In particular, emerging evidence, mainly from animal and *in vitro* studies, suggest a role for kisspeptin on metabolism regulation (for reviews, see Dudek et al., 2018; Wolfe and Hussain, 2018; Hussain et al., 2015). While the metabolic effects of kisspeptins seem to differ between sexes (Tolson et al., 2014), overall the evidence from their involvement in glucose and lipid metabolism suggest that kisspeptins may regulate energy expenditure, with higher levels of kisspeptins related to higher energy expenditure (Dudek et al., 2018; Wolfe and Hussain, 2018; Hussain et al., 2015). Notably, the higher levels of kisspeptins in pubertal boys observed in our study seem in line with known sex differences in metabolism and energy expenditure, known to be lower in females (reviewed in Mauvais-Jarvis, 2015).

We conclude that determining urinary kisspeptin concentrations may be a feasible and sensitive method to quantify circulating levels of these peptides in pubertal populations. The sex differences that were found confirmed results from serum essays in typically developing early adolescents and involved increases in concentrations from early puberty in boys, in contrast with lower and more stable concentrations in girls (except for the transition from pre- to early puberty). Non-invasive methods such as this may aid in the understanding of the differential role of kisspeptins in male and female puberty, such as sex-differences in psychiatric disorders (Worthman et al., 2019), especially considering that kisspeptins seems to modulate various systems that affect behaviors such as mood, fear and anxiety (Mills et al., 2018; Worthman et al., 2019).

Peripheral urinary measurement of kisspeptins can also be helpful in the diagnosis of central precocious puberty, since the concentration of kisspeptins are higher in affected individuals compared to typically developing same-age peers (e.g. Bianco, 2012; Rhie et al., 2011). This condition is much more common in girls (Bianco, 2012) and is possibly related to the effects of kisspeptins in triggering puberty in this sex. In contrast, there is a much higher incidence of delayed puberty in boys (Bianco, 2012), which might be associated with malfunctioning of the mechanisms that lead to the sharp increase of kisspeptins in peripheral tissues as puberty progresses. A comprehensive understanding of the expression, function, and potential molecular mechanisms of kisspeptin/KISS1R in the peripheral reproductive system can also contribute to our general knowledge of typical sexually dimorphic patterns of pubertal development. However, progress in this area will also depend on studies trying to disentangle the reproductive/developmental effects of kisspeptins from their other physiological roles, such as regulation of metabolism, and on investigations of the possible cross-talk between these functions, so that we can gain a deeper understanding of the biological meaning of variations in peripheral kisspeptin concentrations.

Regarding limitations, the cyclicity of the HPG axis pulses may influence variations in kisspeptins in women (Zhai et al., 2017), which were not controlled here. This was also not controlled in any of the prior studies with pubertal samples, so this does not invalidate the comparisons that were made with other studies. The same can be said about the lack of perfect control over the period of the day in which urine samples were collected, which ranged from 10:00 to 16:00h. Had we included more prepubertal girls, or used Tanner staging, it is possible that we would have shown the increase in kisspeptins from this stage to early puberty that was reported by other researchers (e.g. Zhu et al., 2016). On the upside, unlike most of the published literature, we tested a sample from a developing country, with wide variations in SES and BMI, both of which were controlled for and did not affect the urine concentrations of kisspeptins.

## Acknowledgments

This work was supported by the São Paulo Research Foundation (FAPEP: # 2016/14750-0, 2018/06374-3 and 2019/11706-8), as well as CAPES (finance code 001), AFIP and CNPq (#301899/2019-3). The authors thank Luanna Inácio, Beatriz Zappellini, Robson Kojima, Isis A. Segura, Gislaine A. Valverde Zanini, Mariana Libano, Bianca Rodrigues, Eveli Truksinas and Diogo Marques for help with data collection.

